# Why Do Patients with Type 2 Diabetes Lose Less Weight on GLP-1s? A Mathematical Modeling Analysis

**DOI:** 10.64898/2026.09.08.750112

**Authors:** Kevin D. Hall

## Abstract

**Objective:** To investigate the extent to which attenuation of weight loss observed in patients with Type 2 Diabetes (T2D) treated with semaglutide can be quantitatively explained by changes in energy expenditure and urinary glucose excretion (UGE) that occur with improved glycemic control.

**Methods:** A previously validated mathematical model of human energy balance dynamics was modified to simulate patients with T2D by including the effects of hyperglycemia on energy expenditure as well as UGE calculated based on estimated glomerular filtration rate (eGFR), renal threshold (RT) for glucose reabsorption, and estimated plasma glucose derived from measured glycated hemoglobin (HbA1c). Treatment with sodium glucose co-transporter 2 inhibitors (SGLT2i) was simulated by lowering RT.

**Results:** Attenuated semaglutide induced weight losses in T2D patients without background SGLT2i therapy were partially explained by changes in UGE and energy expenditure. Patients with T2D on stable background SGLT2i therapy were predicted to have the greatest attenuation of weight loss.

**Conclusions:** Changes in energy expenditure and UGE associated with glycemic improvements can substantially offset negative energy balance in patients with T2D treated with semaglutide, especially in patients on stable background SGLT2i therapy.

**Study Importance Questions:**

- **What is already known?**: Patients with Type 2 Diabetes (T2D) lose significantly less weight on GLP-1 based pharmaceuticals than patients with obesity alone. The physiological basis for this gap remains poorly understood.
- **What are the new findings?**: Mathematical modeling demonstrates that reduced energy expenditure and urinary glucose excretion (UGE) associated with glycemic improvements during semaglutide treatment can play important roles in attenuating weight loss in people with T2D, especially in those on background therapy using sodium glucose co-transporter 2 (SGLT2) inhibitors.
- **How might results change clinical practice or research?**: These findings may help clinicians interpret attenuated weight loss in patients with T2D and set individualized expectations. The findings may also help pharmaceutical companies and regulators interpret data on weight loss differences between different patient populations and design trials to better understand weight loss variability in patients with T2D.

## Introduction

Randomized controlled trials of GLP-1 based obesity pharmacotherapy in patients with **type 2 diabetes (T2D)** typically result in less weight loss than in trials including patients with obesity but without T2D – an effect has been observed with semaglutide (1), tirzepatide (2, 3), cargrisema (4, 5), and retatrutide (6, 7) treatments. This weight loss gap is potentially clinically meaningful because T2D remission is correlated with the magnitude of weight loss, and more than 80% of patients who lose 15 kg or more after 12 months put T2D into remission (8).

The mechanisms underlying the T2D weight loss gap in patients with obesity treated with GLP-1s are unclear, but energy metabolism differences in the state of T2D provide some plausible hypotheses. In particular, many patients with T2D may have a baseline “energy sink” in the form of **urinary glucose excretion (UGE)** (9) as well as increased energy expenditure (10), both of which correlate positively with glycemia. Because GLP-1 based therapies improve glycemic control in patients with T2D, any baseline UGE energy sink or elevated expenditure may be diminished during treatment and thereby attenuate net negative energy balance as compared to a patient without T2D who reduces energy intake by the same amount.

Previous post hoc analyses of randomized controlled trials of semaglutide (11, 12) and tirzepatide (13) have shown that the magnitude of weight loss is inversely related to baseline HbA1c as would be predicted by glycemic effects on UGE and energy expenditure. There is also evidence for a T2D weight loss gap in patients with other obesity pharmacotherapies, including naltrexone bupropion (14, 15), phentermine topiramate (16), and lorcaserin (17, 18). Bariatric surgery also appears to result in a T2D weight loss gap (19-21) and while early studies of low-calorie diet interventions found that patients with T2D lose less weight (22-24), a systematic review and meta-analysis failed to find a significant difference (25). However, this null effect was primarily due to the outsized influence of a single study (26). Therefore, attenuation of weight loss with T2D may be a general phenomenon.

To help quantitatively understand attenuated weight loss with GLP-1 treatment in T2D, this study mathematically models the potential magnitude of glycemia dependent expenditure and UGE effects on dynamic energy balance and weight change during treatment with semaglutide in patients with T2D as compared to patients with obesity but not T2D in the STEP UP T2D (27) and STEP UP (28) trials, respectively.

## Methods

I first simulated semaglutide treatment in participants without T2D in the STEP UP trial data using the previously described strategy to model how semaglutide affects appetite and energy intake (29). For these simulations, I assumed that there was no effect of glycemic change on energy expenditure since the participants’ baseline glycemic control was in the normal range (average HbA1c was ∼5.7%) and therefore UGE was zero.

To model patients with T2D, I modified my previously validated mathematical model of human energy balance and body composition change (29) to incorporate UGE as a function of glycemic control and renal function using the following equation:

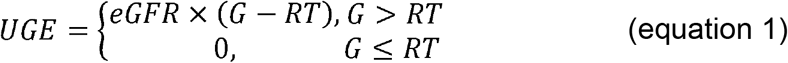

where **eGFR** is the estimated glomerular filtration rate, **G** is the plasma glucose concentration (fasting or postprandial), and **RT** is the renal threshold for glucose reabsorption (30). The STEP UP T2D study reported the mean eGFR = 92 ml/min/1.73 m^2^ which indicated normal renal function, so I assumed a correspondingly normal RT = 180 mg/dl. Body surface area was calculated using the duBois formula (31). For patients with T2D treated with a **sodium-glucose cotransporter 2 inhibitor (SGLT2i)**, I assumed that RT was decreased to 110 mg/dl thereby increasing UGE according to equation 1.

The measured average changes **glycated hemoglobin** (**HbA1c**) reported in clinical trial data were used to estimate average plasma glucose using the **A1c-Derived Average Glucose (ADAG)** study (32). Fasting plasma glucose was also estimated from HbA1c using published regression models (33). The model assumed that fasting glucose prevailed for 8 hours daily, with postprandial glucose concentrations calculated for the remaining 16 hours such that the estimated daily average glucose was equal to the value determined by the ADAG equations.

I simulated semaglutide treatment in patients with T2D to compare with the STEP UP T2D trial data by inputting the measured average time course of HbA1c, but without changing the best-fit model parameters determining how semaglutide affected appetite in the STEP UP study in patients with obesity alone. To model the effects of improved glycemic control to reduce energy expenditure in T2D, I used the estimated fasting plasma glucose values derived from measured HbA1c (33) and decreased energy expenditure by 3.5 kcal/d per mg/dl reduction in fasting plasma glucose (34).

Approximately 30% of the semaglutide treated patients were receiving background SGLT2i therapy in the STEP UP T2D study for at least 90 days and were weight stable (27). Unfortunately, data from the trial were not segmented by background SGLT2i therapy, so I simulated the results with and without SGLT2i therapy which were then averaged according to the 30:70 proportion of STEP UP T2D participants.

## Results

Weight loss in patients with T2D was substantially attenuated as compared to obesity alone (**Figure 1**). In the STEP UP trial involving participants with obesity without T2D, weight loss averaged approximately 18% and 21% at 72 weeks with 2.4 mg and 7.2 mg semaglutide treatment, respectively, whereas the STEP UP T2D trial showed an attenuated weight reduction of approximately 11% and 14% at 72 weeks in patients with T2D.

**Figure 1.**
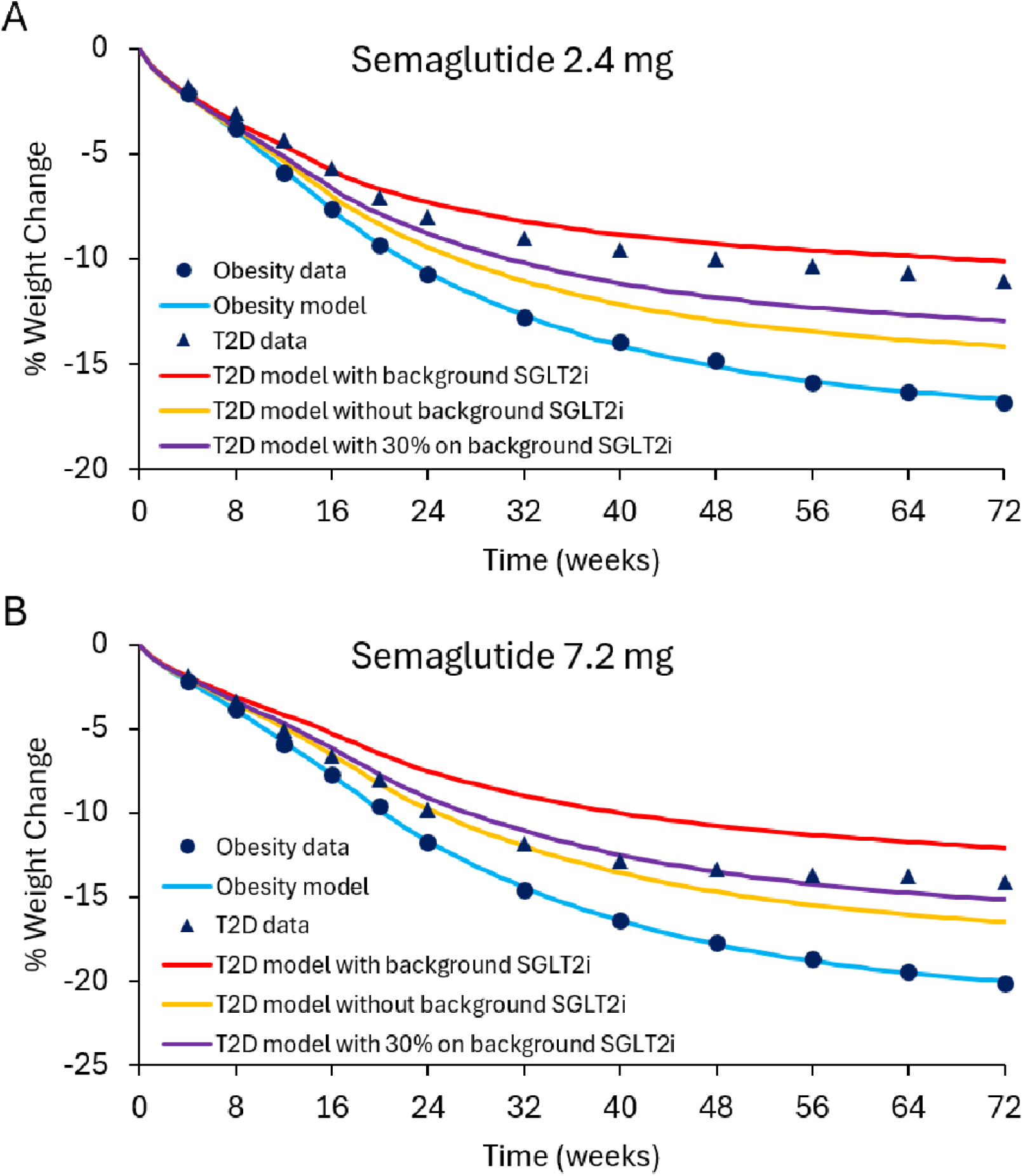
Model-simulated weight loss trajectories during treatment with A) semaglutide 2.4 mg and B) semaglutide 7.2 mg in the STEP UP and STEP UP T2D trials. Model parameters affecting appetite were fit to match the data (circles) in participants with obesity alone (blue). With these appetite parameters fixed, the observed effects of semaglutide on HbA1c were used to estimate changes in glycemia and the effects on energy expenditure and urinary glucose excretion in patients with type 2 diabetes. Background SGLT2i therapy resulted in the greatest attenuation of weight loss (red curve) as compared to simulations of patients with T2D who were not on background SGLT2i (yellow curves). Model simulations with and without background SGLT2i therapy were averaged (purple curves) to represent the STEP UP T2D data (triangles) where 30% of participants were on background SGLT2i therapy.

The mathematical model parameters defining the appetite effect of semaglutide were fit to data in participants with obesity alone (blue curves). Under the assumption of no background SGLT2i therapy, Figure 1 shows that the model simulations (yellow curves) only partially explained the observed weight loss attenuation with T2D. Improved glycemic control during semaglutide treatment resulted in simulated reductions in energy expenditure of ∼175 kcal/d and UGE was reduced to zero from ∼20 g/d at baseline. However, T2D patients on stable background SGLT2i therapy were predicted to have a substantial attenuation of weight loss (red curves) due to UGE reducing from a baseline of ∼125 g/d to ∼40 g/d corresponding to improved glycemia. Averaging the simulation results with and without background SGLT2i therapy in a 30:70 proportion to match the STEP UP T2D study resulted in weight losses (purple curves) that were similar to the observations in the 7.2 mg semaglutide group, but slightly greater than the 2.4 mg group.

## Discussion

Mathematical model simulations indicated that reductions in energy expenditure and urinary glucose excretion associated with improved glycemia likely contribute to the attenuated weight loss with semaglutide treatment in patients with T2D, especially in those with high levels of baseline UGE who substantially improve glycemic control. In particular, the model predicted that background SGLT2i therapy was an important determinant of attenuated weight loss during semaglutide treatment. This result is particularly relevant given that SGLT2i therapy has become part of the standard of care for T2D (35) and an increasing number of patients beginning weight management medications will already be on background SGLT2i therapy.

Given that SGLT2i therapy has been shown to result in weight loss in patients with T2D (36, 37), it may be counterintuitive that background SGLT2i was predicted attenuated weight loss with semaglutide treatment. In the simulations of the STEP UP T2D trial, background SGLT2i was simulated to result in a baseline elevation of UGE by ∼100 g/d as compared to those not on SGLT2i – an amount that depends on model assumptions about glycemia, GFR, and RT according to equation 1. Therefore, on a stable SGLT2i background therapy, introduction of semaglutide results in reduced energy intake and improvements in glycemia that result in large decreases in UGE that offset the negative energy balance as compared to patients who were not previously on background SGLT2i therapy. However, initiation of both SGLT2i and GLP-1 based therapies at the same time is predicted to result in greater weight losses in T2D than either therapy alone.

Unfortunately, the STEP UP T2D study did not report data separately for patients with or without background SGLT2i therapy. Model simulations assumed that glycemic improvements estimated from measured HbA1c changes were similar between these groups. In support of this assumption, previous studies have shown similar reductions in HbA1c in patients with T2D treated with semaglutide regardless of whether they were on background SGLT2i therapy (38, 39). Nevertheless, it would be preferable to model glycemic changes and body weight data separately in patients with and without background SGLT2i therapy.

The real-world impact of reduced energy expenditure and UGE during treatment with GLP-1 based therapies in T2D is likely sensitive to individual variability in energy metabolism, glycemic control, and renal physiology. For example, the magnitude of increased energy expenditure in patients with T2D and the degree to which it changes with glycemia are not well understood but are believed to be related to the energy cost of gluconeogenesis (40). I modeled the energy expenditure effect using data from cross-sectional data (34) and it is encouraging that similar magnitude effects have been observed with longitudinal changes in glycemia occurring during development of T2D and natural weight gain (40, 41). However, it is unclear whether the modeled glycemia related reductions in energy expenditure during semaglutide treatment are accurate.

Another source of variability involves the relationship between HbA1c and average glucose concentrations, especially across different ethnicities (42). I used the standard ADAG equations that are representative of a mostly Caucasian cohort (32) and the estimated average glucose concentrations predicted a relatively modest baseline UGE under assumptions of normal renal function. Other assumptions for the relationship between HbA1c and average glucose concentration, for example using equations derived from the **Diabetes Control and Complications Trial (DCCT)** (43), predicted substantially greater baseline UGE and thereby predict a larger attenuation of weight loss in patients with T2D (not shown). However, the ADAG authors argue that **continuous glucose monitoring** (**CGM**) used in their study, but not in the DCCT, provided a more accurate reflection of average daily glucose (32). A recent study using CGMs highlights the variability of the relationship between average glucose concentrations and HbA1c. While the ADAG equations were likely inapplicable for some ranges of HbA1c in some ethnicities, the results were broadly consistent with ADAG equations in the range of HbA1c changes found in the STEP UP T2D study (42).

Because HbA1c is a lagging indicator of glycemia, the modeled glycemic effects are also somewhat delayed which may result in model simulations with transient inaccuracies in the early periods of GLP-1 based therapy. Future studies utilizing CGM data might provide a more accurate means to assess glycemic parameters for use in the model. Another limitation of these analyses is that the renal threshold for glucose was not directly measured, necessitating the use of literature-derived estimates. Repeated measurements of UGE and energy expenditure could provide model-independent assessments of their contribution to weight loss attenuation in patients with T2D.

A limitation of this study is the reliance on aggregate mean data from the STEP UP and STEP UP T2D trials rather than individual participant data. Using population mean data may obscure significant phenotypic heterogeneity. For instance, there remains substantial uncertainty regarding the magnitude of the UGE and energy expenditure effects and their variability between individuals. Furthermore, differences between the trial populations apart from T2D could also contribute to the observed differences in weight loss with semaglutide treatment. In particular, the proportion of females was higher in STEP UP vs STEP UP T2D (∼74% vs 52%, respectively) and GLP-1 based therapies typically result in greater weight loss in females for reasons that are presently unclear (44, 45).

In conclusion, glycemic dependent reductions in energy expenditure and UGE can contribute to attenuated weight loss T2D. This mathematical modeling study suggests that these effects may partially explain the weight loss gap between patients with and without T2D observed in clinical trials of GLP-1 receptor agonists. Future studies should directly measure dynamic changes in UGE and energy expenditure to test these model predictions. These findings may help pharmaceutical companies and regulators interpret data on weight loss differences between different patient populations and may help clinicians set appropriate expectations for their patients with T2D.

## Acknowledgements

Vian Azzu, Ben Challis, Holly Kimko, Dayna McGill, Jan Oscarsson, and Shivendra Tewari provided helpful comments and suggestions.

## Notes

**FUNDING**: Supported by AstraZeneca Pharmaceuticals.

### Competing Interest Statement

The author is an employee of AstraZeneca Pharmaceuticals.

